# An improved experimental model of cystic hydatid disease in liver resembling natural infection route with stable growing dynamics and immune reaction

**DOI:** 10.1101/155168

**Authors:** Ruiqing Zhang, Xinhua Chen, Hao Wen

## Abstract

Cystic echinococcosis is an endemic parasitic infection in Xinjiang, China and is causing serious economic burdens and public health concerns. An experimental murine model *in vivo* for hepatic cystic echinococcosis was established in C57B/6 mice by injection with human protoscolices via the portal vein of three different concentrations. Mice were followed up 10 months by ultrasound, gross anatomy, pathological and immunological examinations. The protoscolice migration in portal vein, hydatid cyst growth, host immune reaction and hepatic histopathology were examed periodicly. The infection rate of the mice in the high, medium, and low concentration groups were 90%, 100%, and 63.6%, respectively. The protoscolices migrate in the portal vein with blood flow, settle in the liver and develop into orthotopic hepatic hydatid cysts, resembling the natural infection route and course. This study established an improved experimental model of low biohazard risk but stable growing dynamics and immune reaction. It is especially useful for new anti-parasite medication trials agains hydatid disease.

**Summary statement:** An experimental murine model of cystic echinococcosis was set up. This orthotopic model resembles primary infection route and natural infectious course with low biohazard risk and high efficiency.

## Introduction

Hydatid disease caused by *Echinococcus granulosus* is a worldwide zoonosis. It is highly prevalent in Xinjiang, China^1^. Humans are accidentally infected by *Echinococcus granulosus* egg-contaminated areas. Infection causes serious economic, medical, veterinary and public health impact^2^. Animal model plays an important role in the translational study for novel drugs, surgical approaches and vaccine development. An ideal experimental model should orthotopically induce hydatid disease in the most affected organ e.g. Liver. The model should resemble the natural infection route and course with a stable and predictable growth. However, traditional animal models exhibit a biohazard risk when feeding animals with parasitic eggs and induce the parasite cyst in the abdomen cavity as the secondary infection^3^. In human, hydatid disease demonstrates a chronic infectious course and it takes decades for the parasite to settle and grow in the liver^2^. The life cycle includes six stages: (1)The adult Echinococcus granulosus, which is about 3-6 mm in length, resides in the bowel of its definite host; (2)Gravid proglottids release eggs that are passed in the feces; (3)These eggs are then ingested by a suitable intermediate host, including sheep, goat, swine, cattle, horses and camels. The eggs then hatch in the bowels and release oncospheres that penetrate the intestinal wall. These oncospheres then migrate through the circulatory system to various organs of the host. (4)At the organ site, the oncosphere develops into a hydatid cyst. The cyst enlarges gradually, producing protoscolices and daughter cysts that fill the cyst interior; (5)The cyst-containing organs are then ingested by the definite host, causing infection. After ingestion, the protoscolices evaginate, producing protoscolexes; (6)The scolexes of the organisms attach to the intestine of the definite host and develop into adults in 32-80 days. After invading into the GI tract, its life cycle then continues in humans. The eggs then release oncospheres in the small intestine. In liver, oncospheres migrate through the circulatory system and produce hydatid cysts.

To set up such an experimental animal model in order to mimic the natural life circle is expensive and time-consuming. In addition to the time and cost, the biohazard risk also cannot be ignored. Oral feeding with parasitic eggs can cause high infection risk for researchers and requires a high-level biohazard lab to perform the studies^4^^−^^7^. Thus, the development of a highly accurate but low contaminate risk animal model to interpret short-term research results would be beneficial. In this study, we established a mouse model by injecting the portal vein with protoscolices obtained from human hydatid cysts. Ultrasound studies detected cysts within 4 months. The protoscolex migration, hydatid cyst formation, growing dynamics, pathological development and immune reactions were followed up until 10 months.

This study proposes a way to circumvent the many problems linked to a an animal model for hydatid cyst closer to natural infection, i.e. ingestion of oncospheres. Feeding animals Echinococcus eggs in the lab is risky because of biohazard for the lab personnel that can accidentally ingest or inhale the eggs. For this reason, most experimental works are done on the peritoneal injections of protoscolices, which reproduce not the natural route of infection (ingestion of oncospheres) but the natural route of secondary echinococcosis (which is what happens when the contents of a cysts are spilled into the peritoneal cavity). In addition, the diasese model also showed the following benefits: it was not on sheep, nor dog but on mouse1. small rodents so that the expeiment can save labor, cost on big animals; 2. Injection from portal vein instead of feeding from mouth so that biohazard of collecting parasite eggs can be avoided; 3. The model bypasses hatching in the samall intestines so that shortern time and avoid evacuation contamination. With the model, we further follow up and prove injected parasite can steadily grow up into hydatid cyst in liver and steadily stimulate hosts immune reaction.

## Results

### Hydatid cyst develop in mouse liver

In natural infection cycle, the adult parasite worms release eggs from feces and contaminate the environment. Eggs can survial for a year even in the drought and freezing environment and accidentally infect human residence by fece-oral route. In human digestive tract the prasite eggs hatch and releases oncospheres that penetrates the intestine mucosa. They migrate passively through blood in the portal vein to reach liver for final settlement. One oncosphere develops into a hydatid cyst. The hydatid cyst grow up with cyst fluid and infective protoscolices. In our experimental model, by injecting the protoscolices into the protal vein directly,we bypass the contractable egg hatch stage in the intestine, and get the primary hydatid cyst in the liver. The final number of the developed cyst in fact depends on many factors: e.g. space in the liver, the viability of the protoscolex.et al. The freshly collected hydatid cyst (figure 1) is the initial and key factor for the sucessful animal model *in vivo*.

**Fig 1.**
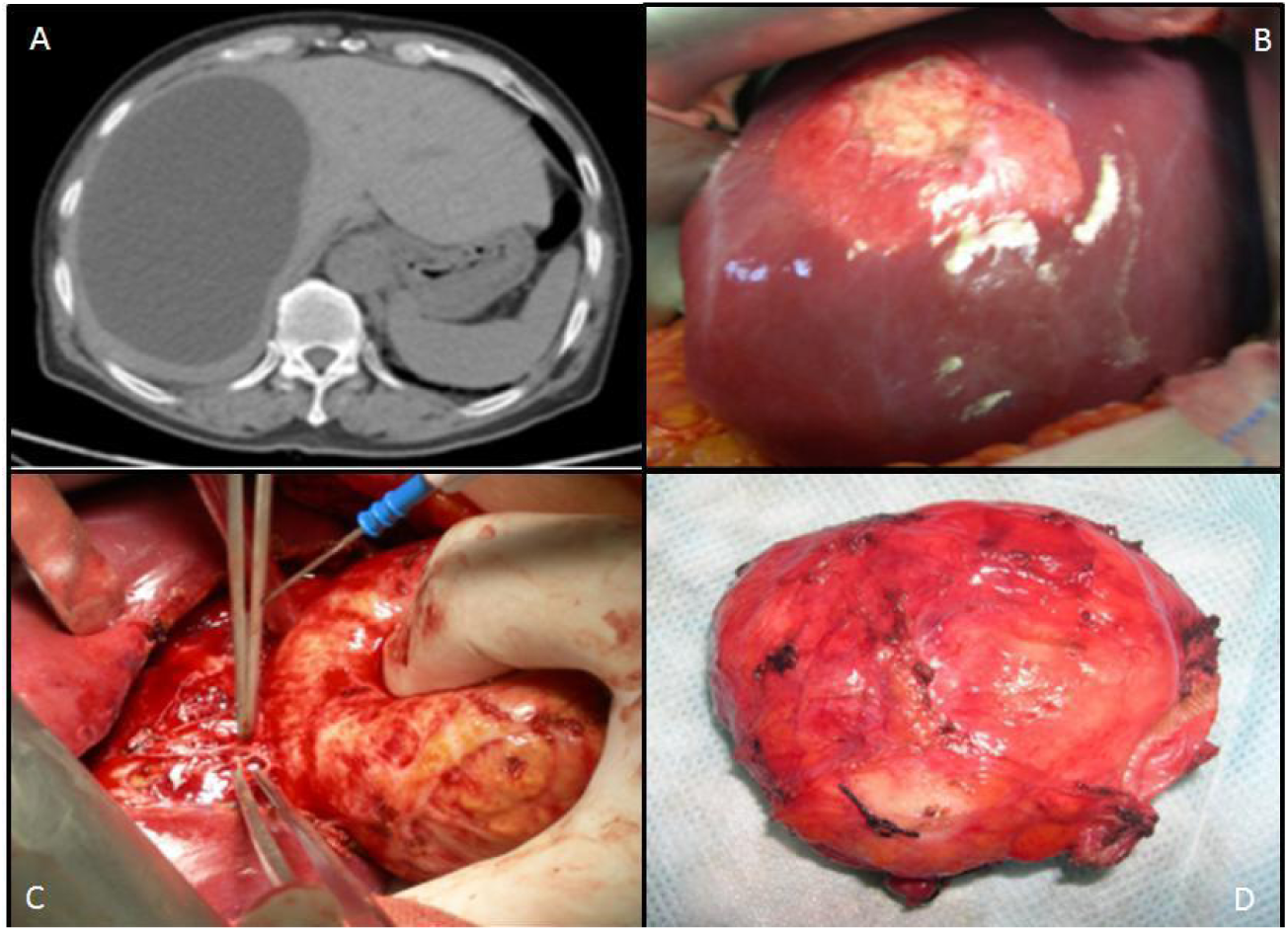
*Echinococcus granulosus* protoscolex collection. The protoscolices were collected from human with hepatic hydatid cyst. WHO classification of Type I hydatid cysts present as a well-defined unilocular and fluid attenuation lesion in liver (figure 1A). The single cyst appearance during an open surgery (figure 1B). the completed removal of the hydatid cyst from liver (figure 1C). The hydatid cyst full of protoscolices inside (figure 1D).

After the mice were injected with protoscolices at three different concentrations (figure 2): (2000/100 μl in Group A, 200/100 μl in Group B and 100/100 μl in Group C), the hydatid disease infection rate of the mice in the three groups were 90% (9/10 in Group A),100% (10/10 in Group B), 63.6% (7/11 in Group C) (Table 1). There was no significant difference among the three groups (P<0.05).

**Fig 2.**
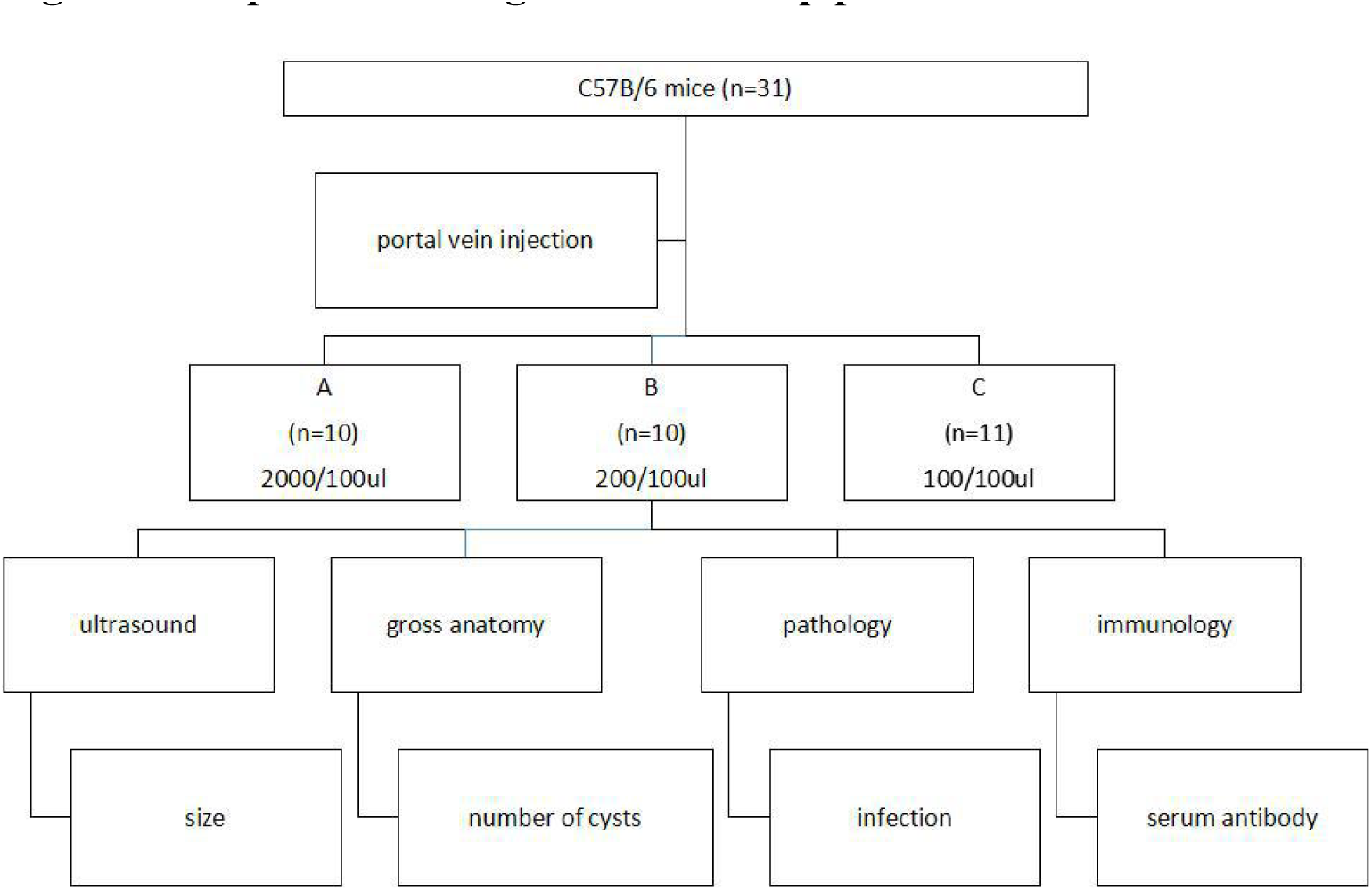
The experiment design and follow up plan. Three different concentrations were prepared for a parallel experiment design with long term follow-up (figure 2). Group A (2000 protoscolices/100 μl), Group B (200 protoscolices/100μl), and Group C (100 protoscolices/100μl). After injection, the hydatid cyst formation, location, distribution and size, pathology and immunology were followed regularly till the 10^th^ month post injection.

**Table 1.** The hydatid disease infection rate of mice in the three groups injected with different concentrations of protoscolices

| Group | Concentration<br>of injected<br>protoscolices | Number of<br>infected mice | Number of<br>non-infected<br>mice | Total | Infection<br>rate (%)* |
| --- | --- | --- | --- | --- | --- |
| Group A | 2000/100 μl | 9 | 1 | 10 | 90.0 |
| Group B | 200/100 μl | 10 | 0 | 10 | 100.0 |
| Group C | 100/100 μl | 7 | 4 | 11 | 63.6 |
| Total |  | 26 | 5 | 31 | 84.55 |

\* $P = 0.096$ , no significant difference in the infection rate among the three groups.

### Hydatid cyst location in the liver

Four weeks after portal vein injection, visual lesions on the liver could be found. After 4 months, the hydatid cysts presented significant growth. Table 2 presents the anatomical locations of the hydatid cysts in the mouse liver (Table 2). Hydatid cysts occurred in any part of the liver and there was no significant preference in any of the liver lobes.

**Table 2.** Lobe position and quantity of hydatid cysts in the mouse liver

| Mouse ID | Group A | Group B | Group C |
| --- | --- | --- | --- |
| 1 | middle lobe: 2 | middle lobe: 2 | right lobe: 3 |
|  | upper right lobe: 1 | lower right lobe: 2 | lateral left lobe: 1 |
|  | lower right lobe: 1 | lateral left lobe: 3 |  |
| 2 | middle lobe: 6 | middle lobe: 1 | lateral left lobe: 2 |
|  | upper right lobe: 5 | lateral left lobe: 1 |  |
| 3 | multiple cysts all over<br>liver | lower right lobe: 1 | none |
| 4 | middle lobe: 6 | upper right lobe: 2 | middle lobe: 3 |
|  | upper right lobe: 2 |  | lateral left lobe: 3 |
|  | lower right lobe: 2 |  |  |
|  | lateral left lobe: 2 |  |  |
| **5** | multiple cysts | middle lobe: 3<br>upper right lobe: 1 | lower right lobe: 1<br>lateral left lobe: 3 |
| **6** | none | lower right lobe: 1 | none |
| **7** | multiple cysts | upper right lobe: 1 | middle lobe: 1 |
| **8** | middle lobe: 1<br>upper lobe: 1 | lateral left lobe: 4 | middle lobe: 2<br>lateral left lobe: 3 |
| **9** | multiple cysts | lateral left lobe: 1 | middle lobe: 1 |
| **10** | multiple cysts | upper right lobe: 1<br>lower right lobe: 2 | none |
| **11** | — | — | none |

The fundamental structure of the four major liver lobes of rat and mouse livers and the segmentation of human liver according to Couinaud is similar and the fundamental structure is comparable. These findings allow the previous use of rodent models in experimental hepatobiliary surgery. The murine and human liver are comparable due to the similarity of the fundamental structures. These findings allow the use of mice to set up the experimental hydatid disease model.

### Pathogenesis and efficiency

After 6 months, when the hydatid cysts were fully developed in the mouse liver, the developed cysts/number of protoscolices injected ratio was evaluated using two markers: pathogenicity (number of cysts per protoscolex) and number of hydatid cysts per mouse. The gross anatomy and column illustration are shown in Fig 3. Cyst abundance in each mouse reflects protoscolex immune reaction, which stimulates the host immune system to produce IgG against the parasite. Although Group A had the highest parthenogenesis (2.395±0.7424) and cyst abundance (47.90±14.848), the condensed lesion made observation of the individual cyst impossible.

**Fig 3.**
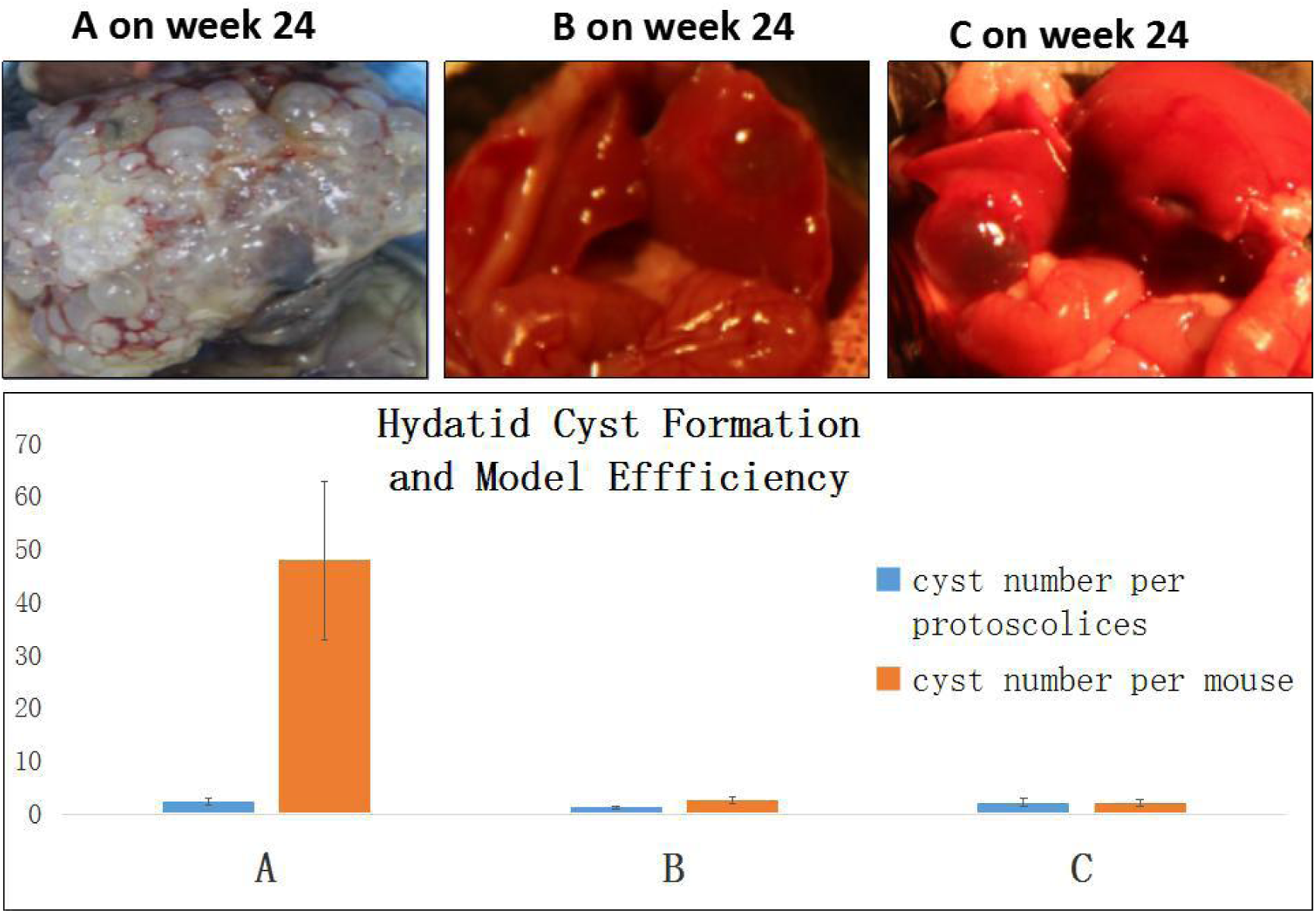
The pathogenesis and efficiency evaluated by cyst per protoscolices. After 6 months when the hydatid cysts fully developed in mouse liver, the injection efficiency was evaluated by two markers: the pathogenicity (cysts number per protoscolex) and hydatid cyst number per mouse.

### Optimization of injection concentration evaluated using a standard score

During the 10-month long follow-up period, no mouse died due to portal vein bleeding, surgery related infection or cachexia, unless a mouse was euthanized during the monthly routine examination. In terms of the hydatid disease model success rate, there was no significant difference among the three groups. However, the experiment model on hydatid requires a more reliable normal distribution. Thus, the standard score was used in this study to compare the reliability and efficiency of the animal models (Fig 4).

**Fig 4.**
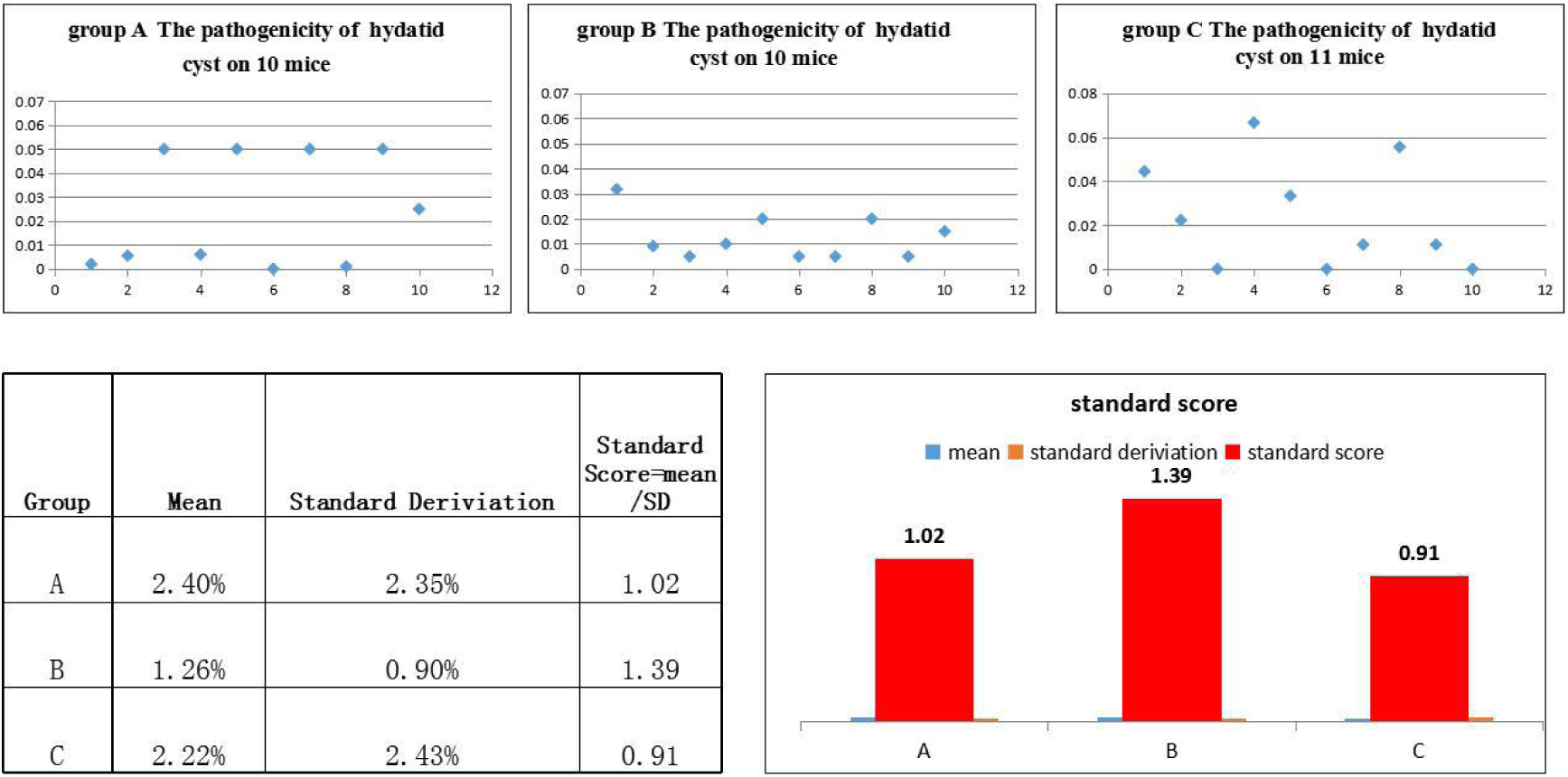
The injection concertation optimization evaluated by standard score. The standard score was used in this study to compare the reliability and efficiency of animal models. The norm distribution of Group A, B, C was shown in Figure 4 upper panel. The standard score was calculated by dividing a mean and a standard deviation (see the calculation in Figure 4). Standard score = (raw score - mean)/SD. A standard score of 1.39 of Group B indicate its performance was better compared with Group A (1.02) and C (0.91).

The standard score was calculated by dividing the mean and standard deviation (see the calculation in Fig 4). This value indicates how well the model reflected the normal distribution compared to other models (the norm distribution of Groups A, B, and C is shown in Fig 4, upper panel). Standard score = (raw score - mean)/SD. This value allows comparisons to be made between the 3 models with different distribution characteristics, i.e., mean and SD. Thus, a score of 1.39 in Group B indicates that its performance was better compared with Groups A (1.02) and C (0.91).

### Migration of protoscolices from the portal vein to liver lobe

The path and course of the protoscolex migration from the portal vein to the liver were tracked by open examination, pathology and ultrasound. On the day of portal vein inoculation with human *Echinococcus granulosus* protoscolices, the branches of the portal vein diameter increased. With congestion of condensed protoscolices (Fig 5, middle panel), 1 day after inoculation, the inflammation cell migration was incarcerated; 3 days after inoculation, a significant inflammatory reaction zone formed; 3 weeks later the protoscolex developed into vesicles (Fig 5, middle panel); and 6 weeks after inoculation, none of the protoscolex could be found but visible vesicular structures of hydatid cyst were observed (Fig 5, middle panel). The open examination showed the distribution and cyst abundance in the livers of Groups A, B and C (Fig 5, upper panel). After 4 months, the ultrasound detected spherical, fibrous rimmed cysts with surrounding host reaction. After 6 months, an even larger parent cyst with satellite daughter cysts within or outside the parent cyst was found (Fig 5, lower panel). The rodents have the four major liver lobes similar to human hepatic Couinaud segments. This murine and human liver are comparable due to the similarity of the fundamental structures. These findings allow the use of mice to set up the experimental hydatid disease model.

**Fig 5.**
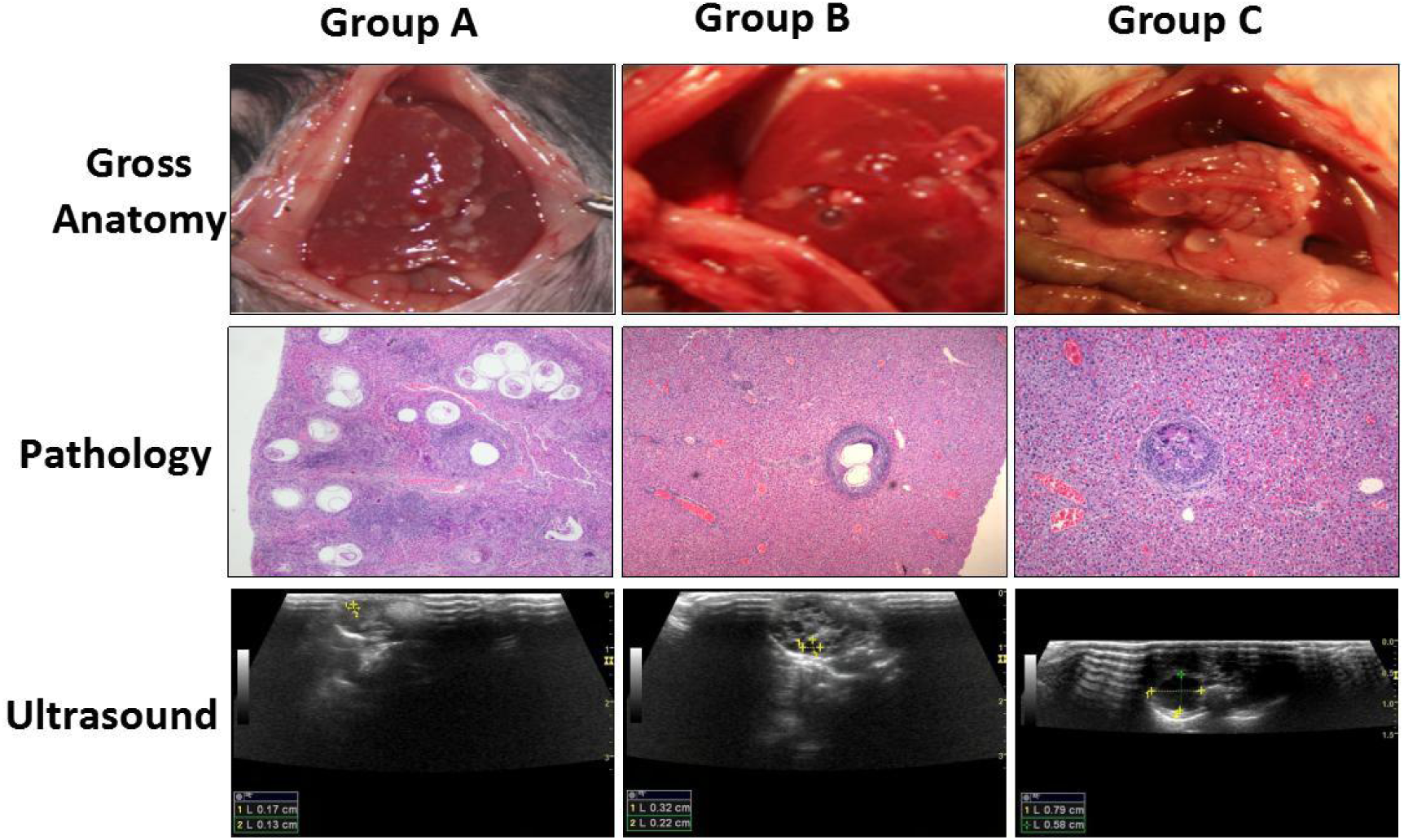
The migration of the protoscolices in the portal vein and liver. The path and course of the protoscolex migration in portal vein and liver were tracked by open check, pathology and ultrasound. When the human Echinococcus granulosus protoscolices were injected they travel from small branches of portal vein to the liver with the blood flow, causing inflammation cell infiltration. When the hydatid cyst formed the infected liver present the infection zone around the parasite lesion. The open check showed the distribution and cyst abundance in liver of Group A, B and C. After 4 months, the ultrasound detected spherical, fibrous rimmed cyst with surrounding host reaction. After 6 months the even larger parent cyst with satellite daughter cysts within or outside the parent cyst were found.

### Hydatid cyst growing dynamics measured by ultrasound

After six months, the ultrasound could detect a stable increase in the number of hydatid cysts (Fig 6). The average cyst diameter in Group B on the 24th, 28th, 29th, 32nd and 36th weeks were 2.48 ±0.91 mm, 3.29±1.86 mm, 3.87±2.26 mm, 5.00±2.57 mm and 7.98±2.75 mm, respectively (Table 3), indicating a significant increase in diameter over time after 6 months (P<0.001).

**Fig 6.**
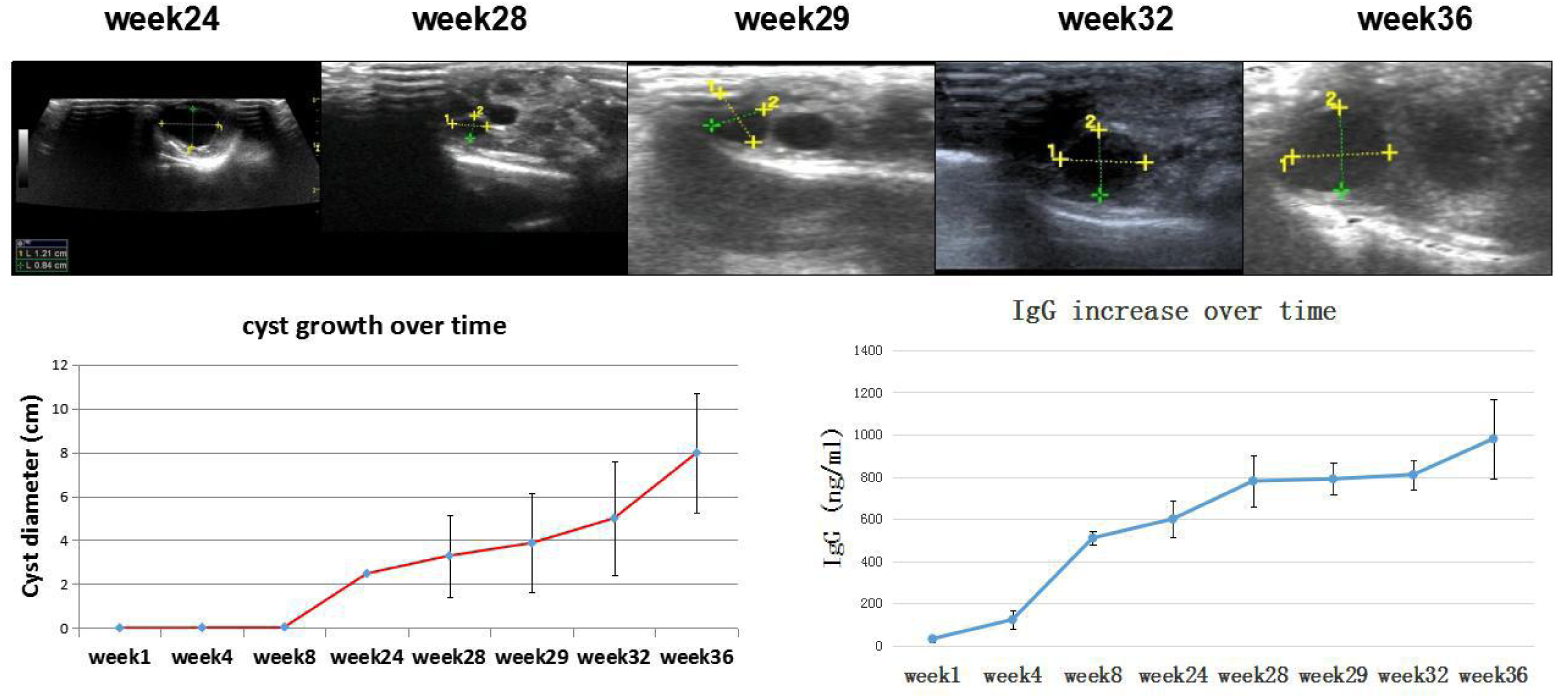
Hydatid cyst growing dynamics measured by ultrasound accompanied by IgG increase. Ultrasound measured the detectable hydatid cysts after 4 months. After six months the hydatid cysts were fully developed and stimulate strong host immune reaction marked by IgG.

**Table 3.** Diameter of the hepatic cystic echinococcosis lesion over time

|  | Week 24 | Week 28 | Week 29 | Week 32 | Week 36 | P |
| --- | --- | --- | --- | --- | --- | --- |
| Cyst diameter<br>(mm) ($\bar{x} \pm S$) | $2.48 \pm 0.91$ | $3.29 \pm 1.86$ | $3.87 \pm 2.26$ | $5.00 \pm 2.57$ | $7.98 \pm 2.75$ | 0.001 |

### Immunological changes evaluated by IgG

When the protoscolices migrated in the portal vein, the host had a low level of IgG (40 ng /ml at week 1). After approximately 6 months, the hydatid cyst became fully developed, and the cyst began to release different antigens to modulate the host immune surveillance. IgG increased in parallel with the hydatid cyst volume (500-800 ng/ml during week 24-week 32). Parasitic antigens stimulated a series of complex host immune responses, which may benefit both the host and parasite for a symbiotic relationship (800-900 ng/ml at week 36) (Fig 6)

### Histopathological changes in the hydatid cyst-infected liver

Microscopic examination of the mouse liver revealed parasitism related pathological changes. After injection, the protoscolices congested the portal veins. They caused dilatation in the vessel sinusoids. Dead protoscolices resulted in focal degeneration and necrosis; and the mouse liver reacted with an increased diameter of central veins. The mouse liver also showed protective immune reactions, such as lymph cell infiltration and fibrosis capsules (Fig 7).

**Fig 7.**
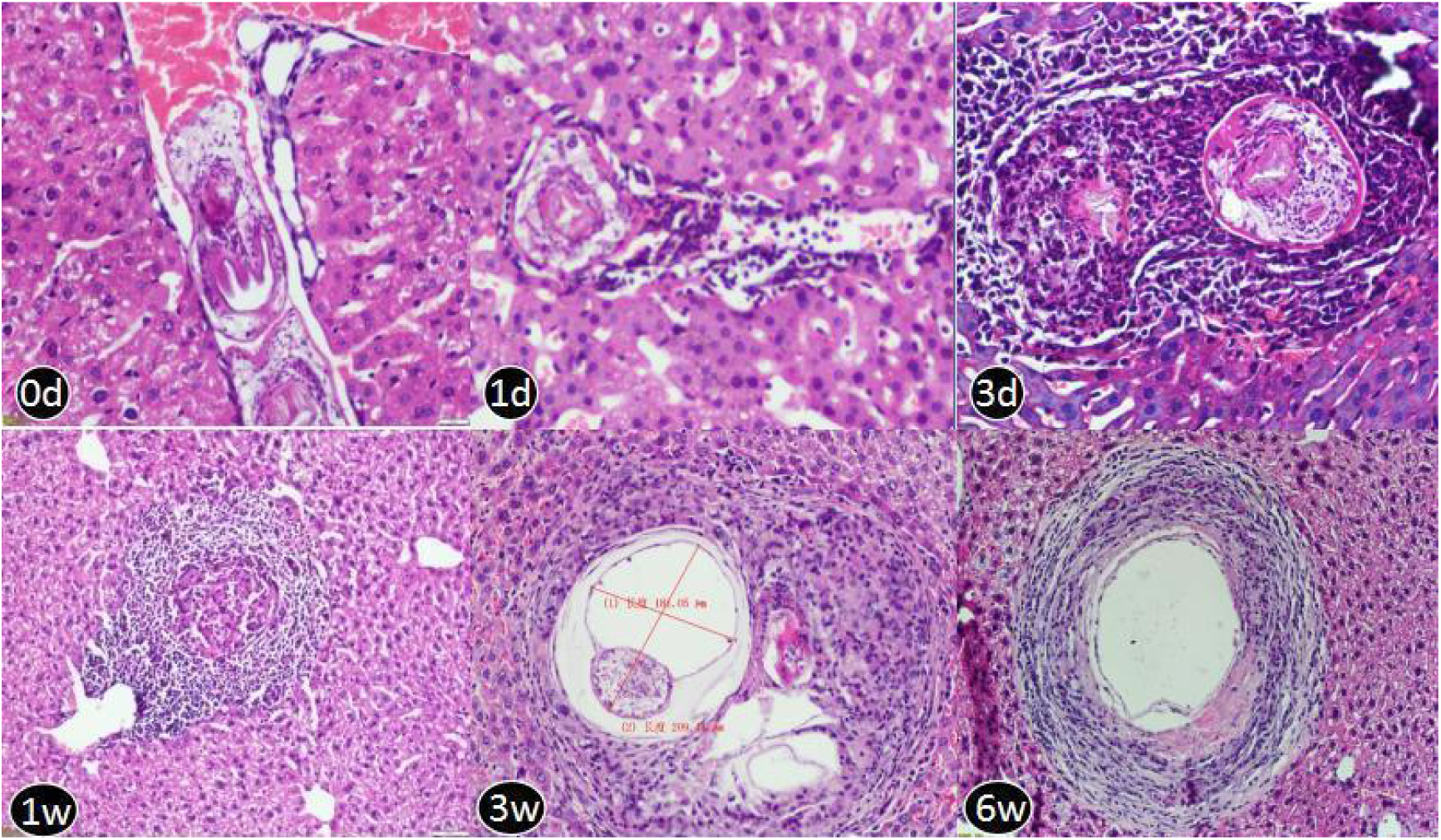
Pathological H&E slides of mouse liver injected with 200 protoscolices at different time points: 0 day,1 day, 3 days,1 week, 3 weeks and 6 weeks. 0 day: On the same day of portal vein injection, the protoscolices congested portal veins. 1 day: The injected protoscolice caused the dilatation in vessel sinusoids. The central vein size increased. 3 day: Those dead protoscolices end up as focal degeneration; mouse liver reacted with increased diameter of central veins and inflammatory cell infiltration (blue stained) and collagen deposition(red stained). 1 week: Mouse liver had protective immune reactions such as lymph cell infiltration. The vessel fibrosis were found in biliary duct and portal vein area 3 week: The newly developed hydatid cyst was found full of germinal layer, laminated layer and the beginning of the adventitious layer. 6 week: the mouse localized the hydatid cyst with compressed and fibrotic host tissue. (H&E stain, 10X)

## Discussion

Hydatid disease is defined as a zoonotic disease or neglected tropical disease. It is a public health problem worldwide. As one of the most serious endemic diseases, it is extremely hazardous in Xinjiang, China due to poor health education and a lack of effective medication^8,9^. Thus, an animal model is needed as the basis for the development of new medication against hydatid disease^10^.

To establish a mouse model of *Echinococcus granulosus*, different infection routes have been investigated, orally, intraperitoneally or intravenously. Different parasitic stages have been employed, such as parenterally with eggs, hatched eggs or activated oncospheres. Generally, less than 1% of the oral dose was established as cysts. In addition to the low infection rate, the generation of experimental animals orally with eggs might pose potential contamination risks to the laboratory personnel who are exposed to faeces of the infected mouse and the liver with hydatid cysts^3,5-7.^

In this study, an effective animal model was established to mimic the natural infection route and course of echinococcosis in human. This animal model showed the following specific advantages.

### Safe operation and low biohazard risk

Echinococcus granulosus poses the greatest risk because it is the most common and widely distributed species. Accidental ingestion of infective eggs is the primary laboratory hazard. A single infective egg from the faeces of the definitive host could potentially result in serious infection. Handling parasites require special care and a special lab facility^4^; in this study, an improved protocol without feeding high risk eggs orally into the gut, but instead injecting protoscolices via the portal vein, reduces the occurrence of laboratory-acquired infection in the laboratory and is safe for animal care personnel.

### Most popular intermediate host

Hydatid disease can use many other wild herbivores as an animal model, such as sheep, goats, cattle, camel, buffalo, swine, and kangaroos, but small rodents showed high feasibility in the general animal centre. A mouse model is easy to perform biochemical exams with mouse-derived antibodies. In this study, the animal model was established using the most popular intermediate host mice. Good susceptibility to human protoscolices and a high yield of hydatid cysts were observed in the liver.

### Natural infectious route and course of orthotopic and primary infection organ (liver)

The hydatid cysts formed by the metacestode (larval stage) migrate from the intestine to the liver via the portal vein and finally develop into hydatid cysts most often in the liver^1,2^. C57B/6 mice were injected via the portal vein with protoscolices from human. This animal model mimics the natural infectious route. Protoscolices migrate from the portal vein into the liver lobe, forming hydatid cysts on the orthotopic and primary infection organ.

### Stable hydatid cyst formation and growth

Small cystic larvae were observed macroscopically in the liver 3 weeks post-injection. The laminated layer was found 6 weeks post-injection. At four months post-injection, larger larval cysts were found in the orthotopic liver. A laminated layer with mature protoscolices was observed to be surrounded by lymphocytes.

### Convinient anatomy location

Mouse liver anatomy is similar to humans except that the human liver has a larger right lobe and a large right portal vein compared with the left side. When the human superior mesenteric vein and splenic vein is confluent in the portal vein, the majority of blood flow goes into the right portal vein, thus carrying more parasites in the flow into the right lobe. The majority of human hepatic hydatid cysts (60% -80%) was found in the right lobe. However, in the mouse model, the liver lobes showed no significant difference in this proportion. The middle lobe exhibited the highest volume percentage (approximately 30% of the total liver volume). Thus, the lesion location in the mouse liver showed no lobe preference and no lobe appeared in the liver. Mice injected with 2000 protoscolices produced more than 100 vesicles by the end of the study (9 months). Mice injected with 200 protoscolices yielded 1-4 vesicles.

Technically, the location of the hydatid lesions could be controlled by fixing the right lobe by selectively blocking the portal vein. To decrease the confounding factors, selective blockade of the portal vein was avoided in this study but it was technically possible in specific circumstances.

### Active hydatid cyst

In human beings, the final fate of chronic hydatid cysts in liver are quite different. Some cysts keep on expanding slowly in decades of years without obvious symptoms. Some cysts grow up to certain volume (e.g. when the cyst’s diameter is larger than 5cm) and become symptomatic. Some cysts rupture spontaneously and the spillage of parasite tissue causes the secondary echinococcosis. Some cysts had necrotic processes leading to a solidification and/or calcification of the cysts. The cysts collapse and gradually disappear. According to ultrasonography features of the hydatid cyst, WHO classified cystic echinococcosis from CE1-CE5:CE1, CE2 are active, CE3 as transitional and CE4 and CE5 are inactive. In this study, the hydatid cysts can form in months (vs many years in human) and the ultrasonography showed the cysts formed in murine livers are CE1, the most active type, making the model a reliable animal model for any further study.

### Detectable host-parasite immune reaction and pathological changes, which are proportional to the expansion of the hydatid cyst

The mice produce the host immune protection following primary injection of protoscolices. Accordingly, the protoscolices developed cyst membranes and capsules that are highly effective in protecting the parasite from host immune destruction. IgG is a marker that reflects the host-parasite immune reaction. When a hydatid cyst develops in the liver, host IgG in the serum is significantly elevated and can be used as an indirect marker for a coarse estimate of hydatid cyst volume and parasitic burden^11^.

### Optimal injection concentration of 200 protoscolices

Group B was optimally injected with a concentration of 200 protoscolices, and the cyst number (2.60±0.618) left sufficient space for intervention and further follow-up observation. In Group B, the number of cysts and protoscolices was proportional to the volume of the cysts. The radius of the individual cyst gradually increased accordingly over time (not linearly). In Group B, 100% of the mice developed hydatid cysts with ultrasound detectable lesions. The hydatid cysts became distended and palpable in 4 months. Group B was superior for research due to its low dose of infection and predictable cyst development as well as better norm distribution. It will benefit experimenters to observe the *in vivo* efficacy of new treatments against hydatid without the need for sacrificing the mouse^12-15^.

In summary, we set up the hydatid disease model not on sheep, not on dog, not on human, but on small rodents so that the experiment can save labor, cost and ethic issue on sheep or human. Inject from portal vein instead of feeding from mouth can avoid collecting parasite eggs with bio-hazard risk. This model can bypass the hatching stage in the intestine so that it saves time and avoid contamination. With the animal model, we further showed the animal model can steadily grow up into hydatid cyst in liver and steadily stimulate host’s specific immune reaction. The proper cyst density and anatomical localization enable accurate monitoring. In this study, larval E. granulosus infection was performed in mice,the most popular experimental intermediate host, was established. Using this experimental model the paraiste cyst growth and immune reaction proportional to the cyst volumecan be examed.

## Material and Methods

### Echinococcus granulosus protoscolex collection

The protoscolices in this study were collected from naturally infected human hydatid cysts during an open surgery in the operation room in the First Affiliated Hospital of Xinjiang Medical University (Fig 1). Informed written consent and an image release agreement were obtained in advance from all subjects. The number of protoscolices was adjusted in 0.9% NaCl solution with a 95% viability rate.

Three different concentrations were prepared for a parallel experiment design with long-term follow-up (Fig 2): Group A (2000 protoscolices/100 μl), Group B (200 protoscolices/100 μl), and Group C (100 protoscolices/100 μl).

### Viability test of Echinococcus granulosus protoscolices

Viability was confirmed using the 1% eosin exclusion test to determine the viability of the protoscolices. The viable protoscolices could exclude the eosin such that they were colourless and mobile, while dead protoscolices stained red. The viability was calculated by the number of viable cells divided by the total number of protoscolices. The protoscolices used for injection had more than 95% viability.

### Mice

Eight-week-old, female C57B/6 mice were purchased from the Shanghai Experimental Animal Center of Chinese Academy of Sciences (Shanghai, China). The mouse weight varied from 20 to 24 g. They were maintained in an SPF level Experimental Animal Center of the First Affiliated Hospital, Xinjiang Medical University and acclimatized in the animal facility for one week before injection.

### Portal vein injection

The animals were shaved, scrubbed, and then moved to a sterile surgical area. The animals were anesthetized with chloral hydrate (300 mg/kg) and remained anesthetized during the operation. A 1.5-cm incision was made from the bladder up to the level of the xiphoid. The skin and muscle layers were retraced by tissue retractors to hold them on the left and right sides. The intestines were carefully moved to one side with sterile gauze to expose the portal vein. The portal vein was located under the pancreas. The needle connected to the syringe filled with protoscolices was inserted into the portal vein and the protoscolex solution was released. After injection, the needle was slowly retracted and a piece of gauze was pressed on the puncture site to prevent backflow of blood for five minutes. The intestines were placed back into the abdomen and the abdominal wall was closed. Mice were maintained on the heating pad for recovery with frequent inspection, and no occurrence of bleeding or infection was found.

### Experiment design and grouping and long-term follow-up after injection

In total, 31 mice were randomly divided into 3 groups (Fig 2):

Group A: 2000 protoscolices in 100 μl saline, via portal vein injection, n=10 mice;

Group B: 200 protoscolices in 100 μl saline, via portal vein injection, n=10 mice;

Group C: 100 protoscolices in 100 μl saline, via portal vein injection, n=11 mice;

After injection, the mice were observed regularly by non-invasive animal ultrasound to measure the cyst formation, location, distribution and size. One mouse from each group was euthanized every month and examined for the presence of cysts.

The liver tissue and hydatid cysts were examined microscopically to record the morphological and pathological changes. The liver and hydatid cyst wall were examined histologically by H&E to track the migration path of the protoscolices from the portal vein to the liver. Blood samples were collected to detect IgG production. The experiment grouping and follow-up design was illustrated using a flowchart in Fig 2.

### Histological examination

Livers and hydatid cysts were fixed in 10% formalin and then embedded in paraffin, cut into 5-μm sections, stained with haematoxylin-eosin, and images were obtained using light microscopy to evaluate the tissue structure and pathological changes.

### Detection of immunoglobulin IgG

Blood samples were collected at different time points for detection of IgG antibodies using a nephelometric technique (Beckman Array 360; Beckman Coulter Instruments, Brea, U.S.A)

### Ethical committee approval

All experimental protocols were approved by the Ethical Committee of the First Affiliated Hospital of Xinjiang Medical University (Approved project number: 20141217003). Informed consent was obtained from all subjects. All methods were performed according to the relevant guidelines and regulations of the Declaration of Helsinki and National Institutes of Health Guide for Care and Use of Laboratory Animals.

### Statistical analysis

SPSS Software 17 for Windows (SPSS Inc., Chicago, USA) was used for the statistical analysis. The differences between groups were determined using t-tests, and P-values less than 0.05 were considered significant. A standard score was used to evaluate the normal distribution of cyst formation efficiency among the three groups.

## Acknowledgments

The authors thank staff from Animal Center, Radiation Department and Pharmaceutical Department for their skilled technical support and care of animals.

